# Chromosome-Level Genome Assembly of Navel orange cv. Gannanzao (*Citrus sinensis* Osbeck cv. Gannanzao)

**DOI:** 10.1101/2023.08.09.552660

**Authors:** Zhiwei Xiong, Hui Yin, Nian Wang, Yuxia Gao

**Affiliations:** National Navel Orange Engineering Research Center, Gannan Normal University, Ganzhou, Jiangxi, China; Citrus Research and Education Center, Department of Microbiology and Cell Science, IFAS, University of Florida, Lake Alfred, FL

**Keywords:** navel orange cv. Gannanzao, navel orange cv. Newhall, genome, fruit ripening-related genes

## Abstract

Navel orange cv. Gannanzao is a variant of the navel orange cv. Newhall (*C. sinensis* Osbeck cv. Newhall) that exhibits an earlier maturation, making it commercially valuable. However, the underlying mechanism underneath its early maturation remains unknown. To address this question, we conducted genome sequencing and de novo assembly of navel orange cv. Gannanzao. The assembled genome sequence is 334.57 Mb in length with a GC content of 31.48%. It comprises 318 contigs (N50 = 3.23 Mb) and 187 scaffolds (N50 = 31.86 Mb). The BUSCO test demonstrates 94.6% completeness. The annotation revealed 23,037 gene models, 164.95 Mb of repetitive sequences, and 2,554 ncRNA. Comparative analysis identified 323 fruit ripening-related genes in navel orange cv. Gannanzao genome, while navel orange cv. Newhall genome contained 345 such genes. These genes were organized into 320 orthologous gene families, with 30.3% of them exhibiting differences in gene copy numbers between the two genomes. Additionally, we identified 15 fruit ripening-related genes that have undergone adaptive evolution, suggesting their potential role in advancing fruit maturation in navel orange cv. Gannanzao. Whole genome sequencing and annotation of navel orange cv. Gannanzao provides a valuable resource to unravel the early maturation mechanism of citrus and enriches the genomic resources for citrus research.

## Introduction

Fruit ripening is a complex physiological and biochemical process that involves chlorophyll degradation, synthesis of flavonoids and carotenoids, aroma development, and fruit softening. Plant hormones, transcription factors, and DNA methylation all play important roles in regulating this process (Bai et al. 2021; Li et al. 2021; Kapoor et al. 2022). As the fruit initiates ripening, developmental signals such as sugars, NO, Ca^2+^, and environmental cues like light lead to the accumulation of reactive oxygen species (ROS), triggering the synthesis and accumulation of abscisic acid (ABA) while concomitantly inhibiting the synthesis and action of gibberellins (GA), indole-3-acetic acid (IAA), and cytokinins (CTK). They also synergistically promote the synthesis and action of ethylene, jasmonic acid (JA), salicylic acid (SA), and brassinosteroids (BR) (Guo et al. 2018; Fernández-Milmanda et al. 2020; Li et al. 2021). Fruits can be categorized into two types: ethylene-dependent respiration climacteric fruits and ethylene-independent non-respiration climacteric fruits (Pech et al. 2008).

Hormones together constitute a complex regulatory network, which orchestrates the maturation of fruits with harmonious precision. The synergistic regulation of ABA, ethylene, and IAA is evident in non-respiratory climacteric fruits (such as citrus). Both ABA-IAA interaction and ethylene-IAA interaction are observed in respiratory climacteric fruits, whereas ABA-ethylene interaction are found in both climacteric and non-climacteric fruits. Those interactions play critical regulatory roles during fruit ripening (Guo et al. 2018; Li et al. 2021; Qiao et al. 2021). In concert with hormonal regulation, transcription factors (e.g., NAC) collaborate with MYB, ethylene, and ABA-related transcription factors to form a regulatory network that responds to endogenous and environmental signals, modulating different aspects of fruit ripening (Li et al. 2018; Cao et al. 2020; Martín-Pizarro et al. 2021; Kou et al. 2021).

Citrus fruits are the largest category of fruits in the world and the third-largest traded agricultural commodity globally, holding a significant position in international agricultural trade and serving as an economic pillar for many citrus-growing regions. However, the limited number of commercially viable citrus varieties and their concentrated ripening periods have hindered the development of the citrus industry. Breeding of early and late-maturing varieties is an important objective in citrus breeding. Navel orange cv. Gannanzao, derived from a bud mutation of navel orange cv. Newhall, is an early-maturing variety known for its early and abundant fruit set, strong resistance, excellent quality, and significant economic impact, making it one of the main cultivated varieties in Gannan, one of the largest naval orange producing regions worldwide (Long et al. 2019). Navel orange cv. Gannanzao ripens at the end of September to early October, whereas other naval orange cultivars ripen around mid-to-late November (Liu et al. 2019). However, the mechanism of its early ripening remains unknown. Here, we sequenced and assembled the chromosome-level haploid genome of navel orange cv. Gannanzao. Additionally, we explored the early ripening mechanism by conducting comparative genomic analysis of navel orange cvs. Gannanzao and Newhall.

## Materials and Methods

### Specimen Collection, DNA Extraction, and Sequencing

Navel orange cv. Gannanzao genome was sequenced in this study. Plant materials were collected from the National Navel Orange Engineering Research Center in Ganzhou, Jiangxi Province, China.

Genomic DNA extraction was performed using the phenol-chloroform method (Rana et al. 2019) and isolated using the Nanobind Plant Nuclei Big DNA Kit (Circulomics inc., Baltimore, MD, USA), following the manufacturer’s instructions. The Illumina library was prepared according to the manufacturer’s protocol (Kozarewa et al. 2009) and sequenced using the Illumina NovaSeq 6000 platform, generating 150-bp paired-end reads. For PacBio sequencing, a HiFi SMRTbell Library was constructed using 15 kb DNA fragments, and sequencing was performed using the Pacbio Sequel II platform. Hi-C libraries were constructed (Belton et al. 2012), which were used for subsequent sequencing with the Illumina HiSeq-2500 platform, generating 125-bp paired-end reads.

### De novo genome assembly and Genome quality assessment

HiFi reads were assembled into scaffolds using Hifiasm v0.15.4 with Overlap-Layout-Consensus algorithm (Cheng et al. 2021). Leveraging the Hi-C sequencing data, the scaffold sequences were then elevated to the near-chromosome level using the LAchesis software (Burton et al. 2013). To attain a chromosome-level genome, manual corrections were made based on the intensity of chromosome interactions, analyzed by the juicebox v1.11.08 tool (Durand et al. 2016). The quality of the assembly was assessed by employing the Benchmarking Universal Single-Copy Orthologs (BUSCO) v5.4.7 (Manni et al. 2021) and the Conserved Core Eukaryotic Gene Mapping Approach (CEGMA) v2 (Parra et al. 2007). Furthermore, the coverage of the assembled genomes was determined by mapping Illumina short reads to the assembly using the Burrows-Wheeler Aligner (BWA) (Li and Durbin 2010).

### Genome Annotation

In our repeat annotation pipeline, we employed a comprehensive approach that combines homology alignment and de novo search to identify repeats throughout the entire genome. Tandem repeats were identified using TRF (Benson 1999) through ab initio prediction. For homolog prediction, the widely used Repbase database was utilized along with the RepeatMasker software (Tempel 2012) and its in-house scripts (RepeatProteinMask), using default parameters to extract repeat regions. In addition, a de novo database of repetitive elements was built using LTR_FINDER (Xu and Wang 2007), RepeatScout (Lian et al. 2015), and RepeatModeler (Flynn et al. 2020) with default parameters. The raw transposable element (TE) library was then constructed, including all repeat sequences with lengths greater than 100 bp and gaps composed of ‘N’ less than 5%. To identify DNA-level repeats, a custom library combining Repbase and the de novo TE library (processed using uclust to generate a non-redundant library) was provided to RepeatMasker.

To predict gene models, we employed a combination of homology-based prediction, ab initio prediction, and RNA-Seq assisted prediction. Homology-based prediction involved aligning genomic sequences to homologous proteins using tblastn v2.2.26 (Camacho et al. 2009) with a threshold of E-value ≤ 1e−5. Gene structure prediction was then performed using GeneWise v2.4.1 software (Birney 2004), based on the matched proteins from reference genomes. For automated de novo gene prediction, we utilized Augustus v3.2.3 (Nachtweide and Stanke 2019), Geneid v1.4 (Blanco et al. 2007), Genescan v1.0 (Burge and Karlin 1997), GlimmerHMM v3.04 (Majoros et al. 2004), and SNAP_2013-11-29. To annotate the genome, transcriptome assembly was conducted using Trinity v2.1.1 (Haas et al. 2013). For the identification of exon regions and splice positions, RNA-Seq reads from leaf, root, and fruit tissues were aligned to the genome using Hisat v2.0.4 (Kim et al. 2019) with default parameters. The alignment results were then used as input for StringTie v1.3.3 (Pertea et al. 2015) with default parameters. Finally, a non-redundant reference gene set was generated by merging genes predicted from the three methods using EvidenceModeler (EVM) v1.1.1 (Haas et al. 2008).

Gene functions were assigned based on the protein sequences’ best match through aligning them with Swiss-Prot (Boutet et al. 2007) using Blastp (Camacho et al. 2009), with a threshold of E-value ≤ 1e−5. To annotate motifs and domains, InterProScan70 v5.31 (Quevillon et al. 2005) was employed to search against various publicly available databases such as ProDom (Servant et al. 2002), PRINTS (Attwood and Beck 1994), Pfam (Mistry et al. 2020), SMRT (Lou et al. 2020), PANTHER (Mi et al. 2019), and PROSITE (Hulo 2006). The corresponding Gene Ontology (GO) (Gene Ontology Consortium 2014) IDs for each gene were assigned according to the relevant InterPro (Hunter et al. 2009) entry.

For predicting protein function, we transferred annotations from the closest BLAST hit in the Swissprot20 (Boutet et al. 2007) database and DIAMOND v0.8.22 (Buchfink et al. 2014) with an E-value <10-5, as well as the NR database (Pruitt et al. 2007) with a similar E-value <10-5. Furthermore, we mapped the gene set to a KEGG pathway (Kanehisa et al. 2017) to identify the most suitable match for each gene.

### Synteny Analysis

To investigate synteny between navel orange cvs. Gannanzao and Newhall, we identified the orthologous genes using OrthoFinder v2.5.4 (Emms and Kelly 2019) with default parameter. According to the structural annotation file (gff3), we extracted the positional in genome and sequence length information orthologous genes. MCScanX software (Wang et al. 2012) with default parameter was performed to complete synteny analysis.

### Selection pressure analysis

Orthologous gene families were identified using OrthoFinder v2.5.4 (Emms and Kelly 2019) with default parameter. Based on functional annotation, we extracted gene families containing genes related to fruit ripening. The nucleotide sequences of each gene family were aligned using the MEGA software and subsequently manually edited. The Ka (nonsynonymous substitution rate) and Ks (synonymous substitution rate) were calculated using the Nei-Gojobori method (Nei and Gojobori, 1986) with the Jukes-Cantor substitution model implemented in DNASP 6 (Rozas et al. 2017). For multigene families, DNASP 6 (Rozas et al. 2017) calculates Ka/Ks for every two sequences and then computes the mean value.

## Results and Discussion

### Genome Sequencing and Assembly

Navel orange cv. Gannanzao was sequenced using PacBio seq ll for long-read sequencing, Illumina NovaSeq 6000 for shot-gun sequencing, and Illumina HiSeq-2500 for Hi-C, which produced 94 Gb (∼267-fold coverage), 48 Gb and 45 Gb data, respectively. Illumina and PacBio data were used to for de novo assembly of the genome. The genome size of navel orange cv. Gannanzao was determined to be 334.67 Mb, with 318 contigs and an N50 value of 3.23 Mb (table 1). Furthermore, there were 187 scaffolds with an N50 of 31.86 Mb. The GC content was 34.48%. With the assistance of Hi-C technology, the genome assembly of navel orange cv. Gannanzao was improved to chromosome level, resulting in 9 pairs of chromosomes. The haploid genome size was 302.61 Mb, accounting for 90.42% of the genome (supplementary table S1, Supplementary Material online). Additionally, 178 scaffolds (32.06 Mb, 9.58%) were unable to be assembled to chromosome level.

**Table 1.**
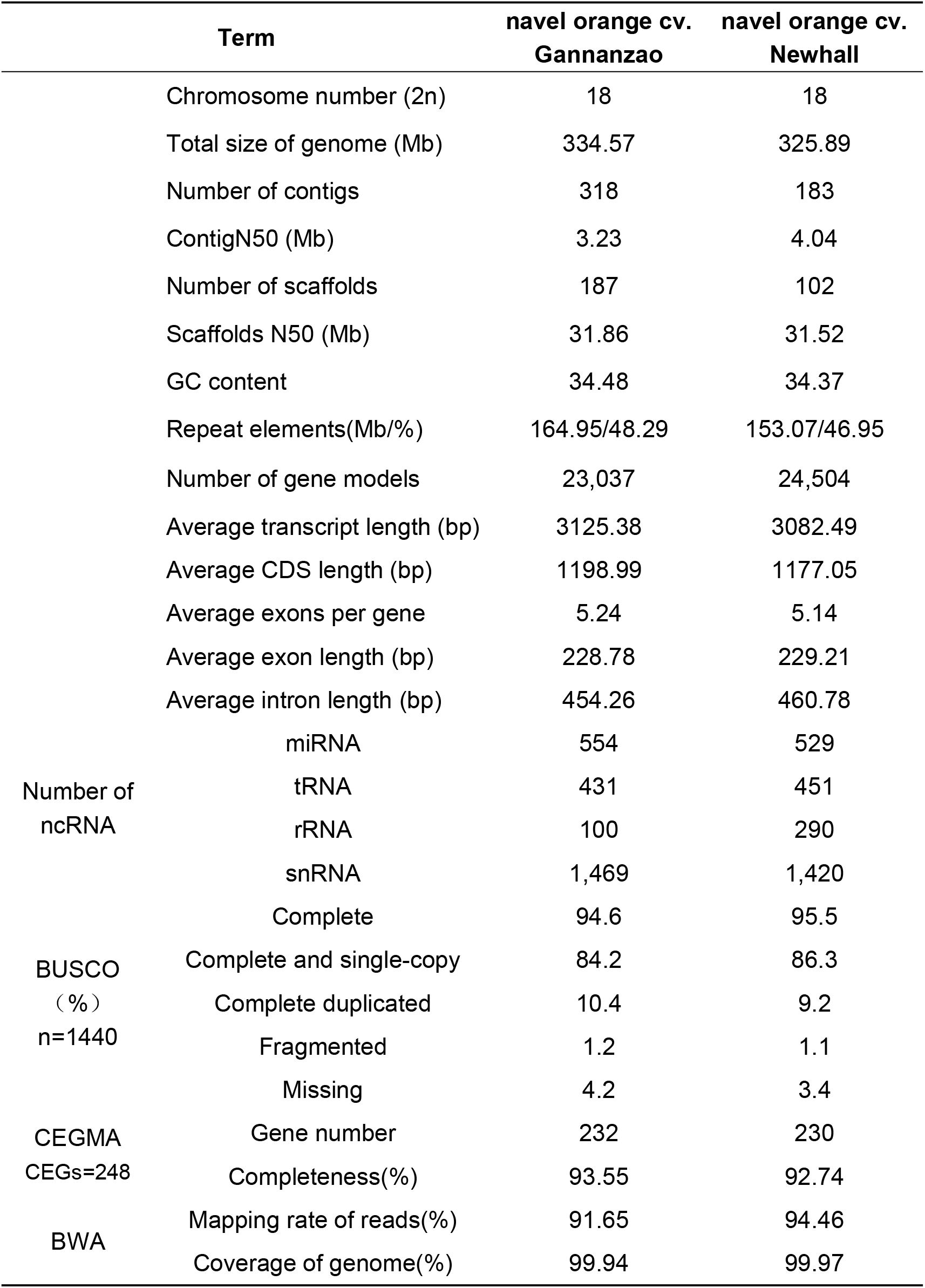
Genome Assembly and Annotation Information of Gannanzao navel orange and Newhall navel orange.

The completeness of the navel orange cv. Gannanzao genome assembly was evaluated using Benchmarking Universal Single-Copy Orthologs (BUSCO) (Manni et al. 2021). The BUSCO completeness was estimated to be 94.6% (table 1), indicating a relatively complete genome assembly. The completeness of the navel orange cv. Gannanzao genome assembly was also evaluated using Core Eukaryotic Genes Mapping Approach (CEGMA) (Parra et al. 2007). This approach utilizes 248 Core Eukaryotic Genes (CEGs) found in 6 eukaryotic model organisms to construct a core gene set. The evaluation revealed that out of the 248 CEGs, 232 were successfully assembled, accounting for a completeness rate of 93.55% (table 1). This result further supports that the genome assembly of navel orange cv. Gannanzao is relatively complete. To evaluate the accuracy of the genome assembly, a small fragment library of reads was selected and aligned to the assembled genome using BWA software (Li and Durbin 2010). Approximately 91.65% of the small fragment reads were aligned with the genome assembly, and the coverage rate was 99.94% (table 1). These results demonstrate a high level of consistency between the reads and the assembled genome. In summary, the genome assembly of navel orange cv. Gannanzao was evaluated using multiple methods, and the results showed high consistency, completeness, and accuracy of the genome.

We compared the navel orange cv. Gannanzao genome sequence with that of navel orange cv. Newhall (Gao et al. 2023) (table 1). The two genomes have similar characteristics, such as genome size, GC content, and values of N50 for contigs and scaffolds (table 1). Taken together, all the data indicate that the genome assembly of navel orange cv. Gannanzao is of high quality.

### Genome Annotation

Repeat sequences in the navel orange cv. Gannanzao genome were identified using a combination of de novo prediction and homology-based alignment. The repeat sequences account for 49.29% of the genome (table 1). The repeat sequences were classified into two types: tandem repeat and interspersed repeat. Through annotation, we found that the tandem repeat sequences consist of 14,446,157 bp, accounting for 4.32%. Meanwhile, the dispersed repeat sequences consist of 150,499,547 bp (44.97%) of the genome, with long terminal repeat (LTR) retrotransposons being the most abundant components, accounting for 127,474,950 bp (38.09%) (supplementary fig. S1, Supplementary Material online).

Gene structure annotation revealed a total of 23,037 genes in the navel orange cv. Gannanzao genome, with an average CDS length of 1,199 bp (table 1). Each gene contains an average of 5.24 exons, and the average length of exons and introns are 228.79 and 454.26, respectively (table 1). Comparative analysis of navel orange cv. Gannanzao and three other closely related species (please list the names here) (Xu et al. 2012; Wu et al. 2014; Gao et al. 2023) demonstrates that navel orange cv. Gannanzao exhibits a notable degree of similarity to the closely related species. This finding supported the high quality of the genome structure annotation of navel orange cv. Gannanzao (supplementary fig. S2, Supplementary Material online).

Alignment of the predicted protein sequences obtained from gene structure annotation with a known protein database revealed that 96.8% of the genes could be functionally annotated (supplementary fig. S3 and table S2, Supplementary Material online). Non-coding RNA (ncRNA) refers to RNA that does not code for proteins, such as rRNA, tRNA, snRNA, and miRNA. These RNAs play important biological functions. Through comparison with a known ncRNA database, 554 miRNAs, 431 tRNAs, 100 rRNAs, and 1469 snRNAs were identified in the navel orange cv. Gannanzao genome (table 1).

### Comparative genomic analysis with navel orange cv. Newhall

We conducted a chromosome synteny analysis between navel orange cv. Gannanzao and its parent variety navel orange cv. Newhall. Similar pattern was observed between the two genomes (fig. 1A). However, significant chromosomal rearrangements were observed. Next, we focused on the early ripening mechanism of navel orange cv. Gannanzao. Based on the annotation information of both genomes, we identified 323 and 345 fruit ripening-related genes in navel orange cvs. Gannanzao and Newhall, respectively. Using the orthoFinder software (Emms and Kelly 2019), 320 fruit ripening-related gene families (95 IAA, 94 Eth, 65 NAC, 20 GTK, 18 ABA, 18 GA, 5 JA, and 5 BR) were identified (fig. 1B). Statistical analysis of the copy numbers of fruit ripening-related genes in each gene family revealed that 223 gene families (69.7%) had similar copy numbers in both genomes. Compared to navel orange cv. Newhall, navel orange cv. Gannanzao exhibited a decrease in gene copy numbers in 40 gene families (12.5%) and an increase in gene copy numbers of 57 gene families (17.8%) (fig. 1D). Analyzing the selective pressure on all fruit ripening-related genes showed that 14 gene families had *Ka/Ks* >1(fig. 1C). These 14 gene families, including 15 ripening-related genes in navel orange cv. Gannanzao (1 ABA, 1 GTK, 3 Eth, 2 GA, 3 IAA, 2 JA, 3 NAC), have likely undergone adaptive evolution, possibly playing a role in the early ripening mechanism of navel orange cv. Gannanzao.

**FIG.1.**
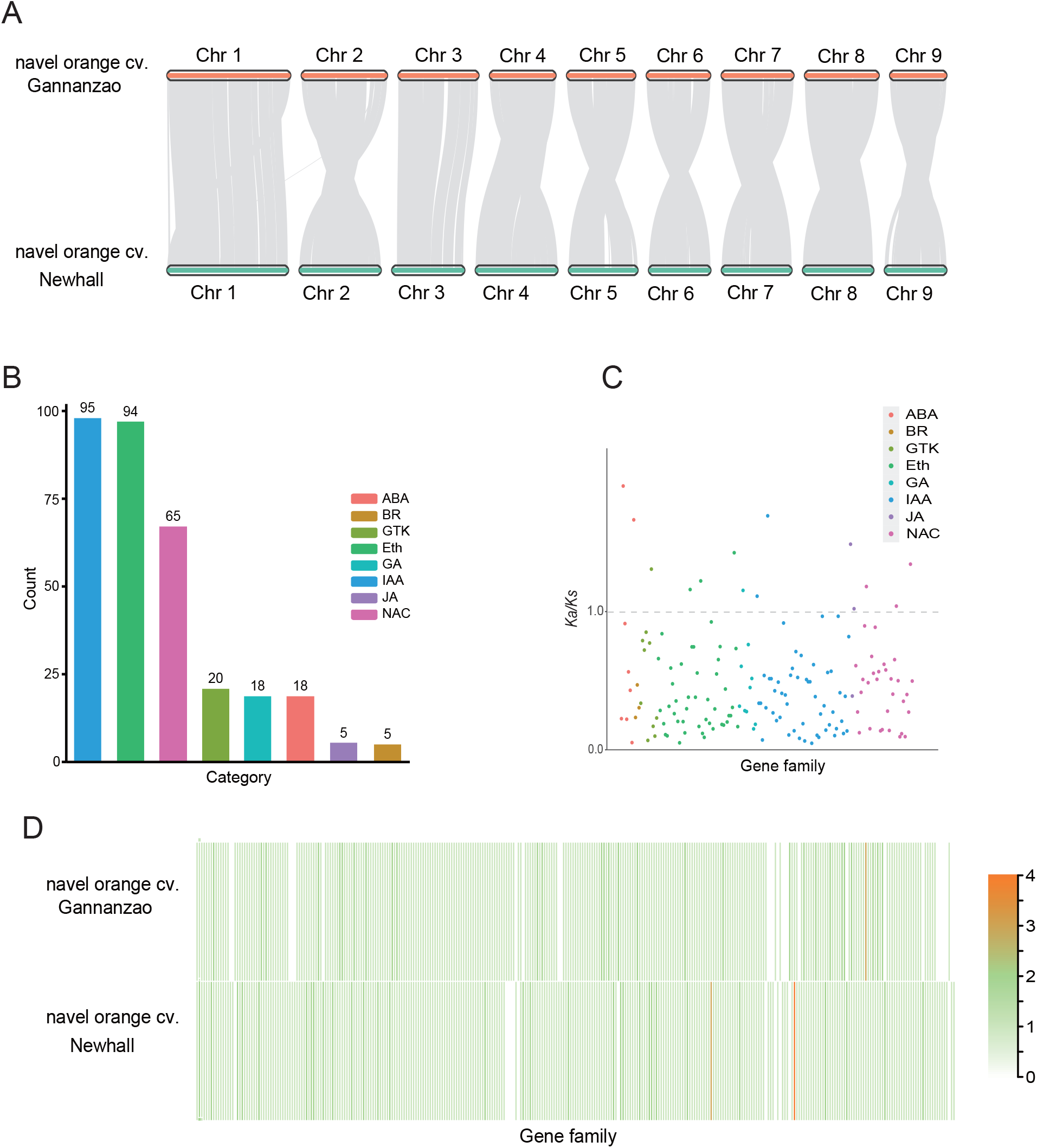
Comparative genomic analysis of navel orange cvs. Gannanzao and Newhall. (*A*) Chromosome synteny plot of two citrus genomes. (*B*) Number of fruit ripening-related genes. (*C*) *Ka/Ks* value of fruit ripening-related genes. The dashed line represents the *Ka/Ks* ratio of 1. (D) The copy number of gene families of fruit ripening-related genes in two genomes.

## Data Availability

The raw sequencing data and genome assemblies of navel orange cvs. Gannanzao and Newhall been deposited in the NCBI databases under BioProject accessions PRJNA997102 and PRJNA810206. The genome annotation is available on the Zenodo data repository under accession doi:10.5281/zenodo.8174988.

## Acknowledgements

ZX and HY collecting and processing data, and generating images, NW contributed technical assistance. Corresponding author YG design of the study, write and submission of the paper, supervised this study throughout and reviewed the article. We thank Guanzhu Han (Nanjing Normal University) for assistance.

